# Organ Chips with integrated multifunctional sensors enable continuous metabolic monitoring at controlled oxygen levels

**DOI:** 10.1101/2024.08.08.606660

**Authors:** Zohreh Izadifar, Berenice Charrez, Micaela Almeida, Stijn Robben, Kanoelani Pilobello, Janet van der Graaf-Mas, Max Benz, Susan L. Marquez, Thomas C. Ferrante, Kostyantyn Shcherbina, Russell Gould, Nina T. LoGrande, Adama M. Sesay, Donald E. Ingber

## Abstract

Despite remarkable advances in Organ-on-a-chip (Organ Chip) microfluidic culture technology, recreating tissue-relevant physiological conditions, such as the region-specific oxygen concentrations, remains a formidable technical challenge, and analysis of tissue functions is commonly carried out using one analytical technique at a time. Here, we describe two-channel Organ Chip microfluidic devices fabricated from polydimethylsiloxane and gas impermeable polycarbonate materials that are integrated with multiple sensors, mounted on a printed circuit board and operated using a commercially available Organ Chip culture instrument. The novelty of this system is that it enables the recreation of physiologically relevant tissue-tissue interfaces and oxygen tension as well as non-invasive continuous measurement of transepithelial electrical resistance, oxygen concentration and pH, combined with simultaneous analysis of cellular metabolic activity (ATP/ADP ratio), cell morphology, and tissue phenotype. We demonstrate the reliable and reproducible functionality of this system in living human Gut and Liver Chip cultures. Changes in tissue barrier function and oxygen tension along with their functional and metabolic responses to chemical stimuli (e.g., calcium chelation, oligomycin) were continuously and noninvasively monitored on-chip for up to 23 days. A physiologically relevant microaerobic microenvironment that supports co-culture of human intestinal cells with living *Lactococcus lactis* bacteria also was demonstrated in the Gut Chip. The integration of multi-functional sensors into Organ Chips provides a robust and scalable platform for the simultaneous, continuous, and non-invasive monitoring of multiple physiological functions that can significantly enhance the comprehensive and reliable evaluation of engineered tissues in Organ Chip models in basic research, preclinical modeling, and drug development.

## 1. Introduction

Organ-on-a-Chip (Organ Chip) technology enables the study of human organ-level physiology and disease states *in vitro*, which has enormous potential for accelerating biomedical research, drug development, and personalized medicine (Ingber 2022). Organ Chips recreate functional, organ-level, tissue-tissue interfaces as they occur *in vivo* by providing structural support and control of chemical signals, mechanical cues, and oxygen levels required for cell and tissue self-assembly. However, despite great progress in this field, more wide-spread use of Organ Chips in research, preclinical, and clinical applications is hampered by technical bottlenecks in creating and maintaining a physiological microenvironment on-chip, which is made more difficult by the lack of non-invasive tools required for assessing multiple microenvironmental conditions and cell metabolic responses simultaneously in real-time.

Current techniques most commonly used for monitoring culture conditions in Organ Chips rely upon use of microscopic imaging of living or fixed chips or “off the chip” analyses of cell viability, proliferation, protein secretion, gene expression, or changes in metabolism using conventional analytical methods (e.g. live/dead stains, cell counting, cytokine analysis, PCR, or various omics techniques) (Ferrari et al. 2020; Fuchs et al. 2021). While useful, these methods only provide minimal information about the local intact tissue microenvironment or dynamic changes in cell metabolism and function over time.

Advances in miniaturized micro-, nano-, and bio-sensor technology have enabled quantitative measurements of a wide range of analytes and microphysiological information (Baracu and Dinu Gugoasa 2021; Chauhan et al. 2021; Pfeiffer et al. 2017). However, sensor integration into Organ Chips remains challenging as most of these systems have complicated designs that are incompatible with multiplexing and automation or are unable to physiologically recapitulate tissue and organ-level structures and functions (Fuchs et al. 2021). In addition, most sensor-integrated chips have been designed for use by the investigator who developed the technology and are not easily adapted for use at a larger scale by multiple laboratories. In addition, while most Organ Chip studies are carried out under ambient air conditions (21% oxygen/ 5% CO_2_), the partial oxygen pressure under normal physiological conditions in most tissues and organs of the body (physioxia) is much lower (4.5-7% oxygen) and these low oxygen conditions influence the physiology and pathophysiology of both host cells and adjacent microbiome (Gan and Ooi 2020). In the most commonly used microfluidic devices that are composed of polydimethylsiloxane (PDMS), recreating physioxia is especially challenging due to the high gas permeability of the PDMS material and the constant exposure to ambient air.

Various methods have been used to recreate physiologically relevant oxygen levels in Organ Chips, including perfusion of anoxic medium through the one channel of a two-channel chip (Shin et al. 2019), culturing chips in anaerobic chambers while perfusing oxygenated medium through one of two channels (Jalili-Firoozinezhad et al. 2019), or wrapping portions of the PDMS chips in low gas permeable films (Grant et al. 2022). All these methods are labor-intensive, technically challenging, and often insufficient to maintain stable low oxygen levels throughout long-term cultures.

To tackle these challenges, we set out to develop Organ Chips made of materials that can enable stable control of low oxygen conditions which also incorporate multiple built-in sensing capabilities for non-invasive, real-time, multifunctional evaluation of the tissues cultured on-chip. Here, we describe the development and validation of a multifunctional sensor-integrated Organ Chip system that employs two-channel Organ Chip devices composed of both highly gas permeable PDMS and low permeable polycarbonate (PC) plastic into which multiple analytical (electrical and chemical) sensors and molecular reporters of cell metabolic activity (ATP/ADP ratio, viability) have been integrated. The multifunctional sensor integrated chip (siChip) is mounted on a printed circuit board (PCB) and designed to fit into a commercially available Organ Chip culture assembly (Pod chip holder; Emulate Inc.) to create a compact and versatile design that can be used as a standalone or linked with other Organ Chip models using a commercially available Organ Chip culture instrument (Zoe instrument; Emulate Inc.). The design of this integrated Organ siChip system allows for easy scale up of manufacturing and handling, automating the operation, and providing remote noninvasive continuous measurements throughout extended culture times. Here, we demonstrate the functionality and robustness of this system using human Gut and Liver siChip models, which maintain oxygen levels in the physiological range (5-7%) on-chip only with cellular respiration and inhibition of oxygen permeability through the PC walls of the chip, and hence, without requiring any user-induced modulation of oxygen levels in the culture medium or incubator. Oxygen levels, pH, transepithelial electrical resistance (TEER), and cell metabolism (ATP/ADP ratio) were also monitored continuously over time in the living chip cultures under baseline conditions and in response to different chemical stimuli (e.g., oligomycin, calcium chelator) as well as co-culture with living *Lactococcus lactis* bacteria.

## 2. Materials and Methods

### 2.1. Design and fabrication of the sensor-integrated chip (siChip)

The microfluidic design of the siChip replicates the configuration of the commercial dual-channel Organ Chip device (Chip-S1; Emulate Inc. USA) to allow its incorporation with a portable module reservoir (Pod™; Emulate Inc.) that contains chambers for storing and collecting the chip media and effluents, respectively; this is accomplished by incorporating a microfluidic compartment that seamlessly connects the reservoirs to the chip enabling automated culture using a commercial culture module (Zoë^TM^ Culture Module; Emulate Inc.) (**Supplementary Fig. S1A,B**). Like the Chip-S1, the dual-channel microfluidic siChip contains two parallel microchannels (1 mm wide × 1 mm high apical channel and a 1 mm wide × 0.2 mm high basal channel) separated by a porous membrane (50 μm thickness, 7 μm pores with 40 μm spacing) (Huh et al. 2013) for co-culturing two cell types, recreating tissue-tissue interfaces, and forming air-liquid or liquid-liquid flow interfaces on-chip. Hollow chambers are located on both sides of the dual channel to which cyclic vacuum can be applied to physically distort the porous membrane and attached tissue-tissue interface and thereby mimic physiological mechanical deformations experienced within whole living organs *in vivo* (e.g., due to breathing, peristalsis, blood pressure pulses, etc.) (**Supplementary Fig. S1A**).

The dual-channel siChip was fabricated by assembling five different layers: apical and basal PDMS layers; apical and basal PC walls patterned with electrodes (for TEER measurements) and oxygen sensors (for oxygen tension measurements); and a PDMS porous membrane that separates the apical and basal channels on which different tissue-derived cells are cultured on its top and bottom (**Supplementary Fig. S1C**).

#### Fabrication of the apical and basal PDMS layers

The apical and basal channel layers were made by first spin coating and curing a single (200 μm, basal channel) or multiple (1000 μm, apical channel) layers of PDMS elastomer (10:1, base:curing agent, Ellsworth Adhesives, Cat. no. 4019862) on PC substrate and curing for 2 hours to overnight at 60°C. The microfluidic features of the apical and basal channels were then made by cutting the designs of the main channels, vacuum chambers, and the inlet and outlet ports in the PDMS layers using a Graphtec (CE5000-60) Vinyl Cutter.

#### PC layers with integrated TEER and oxygen sensors

In-house designed PC sheets with patterned gold electrodes and cut-out inlet and outlet ports (LasX Industried Inc.) were first cleaned with 100% isopropyl alcohol (IPA) and plasma treated on the surface (40 s of oxygen supply at 0.8 mbar and 20% power, Nano-plasma system, Diener electronics GmbH & Co. KG) followed by incubation in freshly made 5% (3-Aminopropyl) triethoxysilane (Sigma-Aldrich, Cat. no. 440140) in deionized water for 20 minutes. Similarly, the apical and basal PDMS parts were plasma treated and incubated in freshly made 1% (3-glycidyloxypropyl)trimethoxysilane (Sigma-Aldrich, Cat. no. 440167) in deionized water for 20 minutes. The silane treated parts were then thoroughly washed with deionized water to remove the excess chemicals, dried with compressed air, and immediately aligned and bonded with the corresponding apical or basal PDMS layers with the gold electrodes facing towards the apical and basal channels. The bonded parts were left at room temperature for 30-40 min to end the silane activation time before their overnight incubation at 60°C under 0.5-1 kg weights.

Oxygen sensors were added to the PC walls of the apical and basal part by depositing small volumes (0.4-0.5 μl) of 10 mg/ml oxygen sensing nanoparticles (OXNANO, Pyroscience) in chloroform slurry at two different locations in each of the apical and basal channels, one location close to the inlet and one close to the outlet ports (**Supplementary Fig. S1C)**. The oxygen nanoparticle sensors were allowed to dry before proceeding to the next step.

#### Fabrication of permeable PDMS membrane

The permeable PDMS membranes separating the apical and basal channels were fabricated in-house using an Automated Membrane Fabricator (AMF 9000) machine. The membrane was designed to be a thin (50 μm) PDMS layer consisting of an array of uniformly distributed pores (7 μm diameter, 40 μm spacing) that allows nutrients and small molecules to travel across the channels but not the mammalian cells. Silicon wafers (4 cm × 4 cm) patterned with an array of pillars (7 μm diameter, 50μm height, 40 μm spacing) were used as a mold to shape the PDMS into the thin, porous membrane. To make these membranes, the wafers were placed in designated slots of the AFM 9000 tray, and PDMS elastomer mix (10:1 base:curing agent) was freshly made and poured onto the center of each membrane wafer post array (0.09 ml/wafer) and allowed to rest for at least 5 minutes to cover >75% of the posts. Square cut pieces (4.5 cm × 4.5 cm) of PC sheets treated with plasma were gently laid onto the PDMS on the plasma-treated side. Each assembly was then layered with a square shape spacer (1 cm PDMS block) on top of the PC sheet for making a uniform membrane thickness. The membranes were placed into the AFM 9000 machine to cure overnight following a stepwise temperature increase from 20 to 60°C at 40 Psi pressure. The next day, PC sheets with attached membranes were gently peeled off the silicon wafer and stored.

#### siChip assembly

To assemble the dual-channel siChip, the apical part and PDMS membrane were first treated with plasma on their exposed surfaces and gently bonded together followed by 2 hours of incubation at 60°C. The PC backing of the membrane was then gently removed and the basal ports were opened using fine-tip forceps to give access to the basal channel. The apical part with the attached membrane and the basal part were then treated with plasma, carefully aligned, and bonded to make the complete dual-channel siChip (**Supplementary Fig. S1C)**. The siChips were then incubated overnight at 60°C in between two flat metal plates with a uniform load of 1 kg weights arrays to ensure full bonding of the apical and basal parts.

#### siChip assembly on a customized PCB

A customized PCB (cPCB, OSH Park, USA) was designed to fit the siChip within the board for direct access to the TEER sensors on-chip and to make a flushed 1.65 mm thick PCB layer that is seamlessly integrated with the Pod^TM^ (Emulate Inc.). The siChip was mounted on the cPCB using the four pads connecting the TEER electrodes to the PCB circuit using silver epoxy conductive adhesive (AI TECHNOLOGY INC, Cat. no. EG8050) (**Supplementary Fig. S1D**) cured overnight at 80°C. To seal-off the integration areas on the cPBC, the top and bottom surfaces of cPCB with incorporated siChip were covered and laminated with a thermal adhesive layer (3M Thermal Bonding Film 583) followed by attaching a 1 mm thick plasma treated PDMS layer (with cut out through-holes to access the channels outlets and oxygen sensors) on top using thermal lamination at 120-140°C (**Supplementary Fig. S1E**).

#### pH sensor integration

pH sensor spots (PyroScience, PHSP5-PK5) were designed to be integrated into the effluent reservoirs of the commercially available Pod^TM^. Frosted surfaces of the reservoirs were coated with a thin layer of silicone glue (SPGLUE, PyroScience GmbH) to maximize the surfaces optical clarity and signal transmission during measurements. The pH sensor spots were then glued to the inner walls of the effluent reservoirs of the Pod^TM^ at the closest proximity to the outlets and allowed to dry overnight in the dark.

#### Modified Pod^TM^ fabrication and assembly

Using Fortus 400mc 3D Printer (ABS-M30 material, Stratasys, USA), we fabricated a base compartment for modified Pod (mPod) to connect the siChip-on-cPCB seamlessly to Pod^TM^ reservoir. The base is constituted of detachable back and front parts with two openings in the back for accessing the cPCB and optical imaging of the culture on-chip, respectively (**Supplementary Fig. S1F**). The chip was assembled on mPod by first placing the siChip-on-cPCB on top of the 3D printed base followed by adding the Pod^TM^ portable reservoir, closing the front insert of the base and then securing all the compartments together using 3 metal screws at the bottom of the base.

#### Modified Tray

A modified Tray (mTray) inspired by the commercial Zoë^TM^ Culture Tray (Emulate, Inc.) was 3D printed using Fortus 400mc 3D Printer (ABS-M30 material, Stratasys, USA) and equipped with electrical connections to connect the pins from the Tray slots to a serial connector located at one end of the mTray for access from outside the incubator via a serial cable. All of the components of the siChip, cPCB, mPod, and mTray were designed using SOLIDWORKS® (Solidworks Premium 2020 SP4.0).

#### Compartment sterilization

All in-house fabricated parts including the siChip, cPCB, and mPod were sterilized using plasma cleaning treatment at 4 s of oxygen supply, 0.3 mbar, and 100% power (Nano-plasma system, Diener electronics GmbH & Co. KG) before chip activation and cell seeding.

### 2.2. siChip activation, cell seeding, and culture

#### siChip activation and extracellular matrix (ECM) coating

The sterilized siChip was desiccated for 30 minutes before 70% ethanol was washed through the apical and basal channels followed by a one-time wash with tissue culture-grade water. The polymeric surfaces of the membrane were then chemically functionalized with 0.5 mg/ml ER1 in ER2 buffer (Emulate^TM^ Inc., USA) under an ultraviolet lamp (Nailstar, NS-01-US) first for 10 minutes followed by additional 5 minutes after refreshing the ER1/ER2 buffer in the channels. Both apical and basal channels were then sequentially washed with ER2 buffer and cold Dulbecco’s Phosphate-Buffered Saline (DPBS). For the Gut Chip, the PDMS membrane surfaces in the apical and basal channels were coated with 30 μg/ml Collagen I (Advanced BioMatrix, Cat. no. 5005) and 100 μg/ml Matrigel (BD Biosciences) in cold PBS. For the Liver Chip, following previously published protocols (Farooqi et al. 2021; Kulsharova et al. 2021; Menon et al. 2014), the apical channel surfaces were coated with 30 µg/ml fibronectin (Corning, Cat. no. 356008) and 400 µg/ml Collagen I (Advanced BioMatrix, Cat. no. 5005) in cold DPBS, while the basal channel was coated with 100 µg/ml Matrigel (BD Biosciences) and 200 µg/ml type I collagen in cold DPBS. The ECM loaded siChips were incubated for 1.5-2 hours at 37°C, followed by one time wash with warm respective culture medium in each channel. Culture medium for the 4 cell lines were vacuum filtered prior to use, to get rid of micro-air bubbles that could hamper the flow in the siChip.

#### Cell seeding

To create the Gut Chip, primary human umbilical vascular endothelial cells (HUVECs; 6-8×10^6^ cells/ml) were first seeded on the basal side of the PDMS membrane and then the chip was immediately inverted and incubated for 2-3 hours at 37°C to allow cell attachment on the basal side of the membrane. The chips were flipped again and the channels were perfused with warm culture medium both apically and basally before seeding human Caco-2 intestinal epithelial cells (Harvard Digestive Disease Center; 2-3×10^6^ cells/ml) in the apical channel. The chips were then incubated overnight at 37°C under static conditions to allow cell attachment on the upper side of the membrane. The next day the chips were perfused with endothelial growth medium (Lonza, EGM^TM^-2 BulletKit^TM^, Cat. no. CC-3162) in the basal channel and Caco-2 growth medium (15% FBS in DMEM) in the apical channel.

To create the Liver Chip, human liver endothelial cells (LECs) (Lonza HLECP1; 4-5×10^6^ cells/ml) were first loaded in the basal channel followed by seeding of human Huh7 hepatocytes (Sekisui JCRB0403-P; 2.5-3.5×10^6^ cells/ml) in the apical channel following the same method described above for the Gut Chip. After overnight incubation at 37°C, the channels were perfused with the Huh7 growth media (10% FBS in DMED) apically and LECs growth media (Lonza, CC-3162) basally before starting the culture.

#### Culture of siChips in the commercially available culture instrument

To culture the seeded siChips in the Zoë^TM^ Culture Module (Emulate, Inc), the Pods^TM^ were first primed with the endothelial and epithelial media in the basal and apical channels, respectively, to remove any air from the microfluidics that connect the reservoirs to the chip. The Pod^TM^ reservoir with the attached microfluidic compartment was then removed from the commercial base, integrated with the siChip on the mPod, and placed in the mTray inside the Zoë^TM^ Culture Module before starting the automated flow perfusion while placed in a 37°C incubator (**Supplementary Fig. S2A**). The Gut siChips were maintained under a continuous flow of Caco-2 growth medium apically and endothelial growth medium basally at 40 μl/hr for 10 days before changing to differentiation-promoting media (5% FBS in DMED apically and 0.5% FBS basally) for the rest of the culture time. The Liver siChips were also maintained under a continuous flow of Huh7 growth medium apically and LECs growth media basally at 30μl/hr for 5-6 days before reducing the media FBS content for induction of tissue differentiation (5% FBS in DMEM, apically and 0.5% FBS, basally).

### 2.3. Continuous measurement of TEER, oxygen, and pH on-siChip

#### TEER measurement and analysis

TEER was measured continuously while siChips were in culture in the mTray inside the Zoë^TM^ Culture. The mTray was connected to a CompactStat.h Mobile electrochemical interface (B32121, Ivium Technologies B. V., The Netherlands) outside the incubator using a serial cable and the TEER was measured following the method previously reported by Henry et al. (Henry et al. 2017). Briefly, a four-point impedance measurement was taken periodically every 1-2 days using the IviumSoft application (V4.97, Informer Technologies, Inc.) paired with the CompartStat h electrochemical interface. A current of 100 μA was applied between the two excitation electrodes located on the opposite sides of the siChip membrane and the drop in the potential was measured in the second set of electrodes. For each chip, the impedance spectra were recorded over a range of frequencies varying from 10Hz to 100kHz.

The impedance at high frequencies (>10kHz) is mainly characterized by the media resistance while at lower frequencies (<100Hz) the TEER from the tissue electrical resistance dominates the impedance signal (Henry et al. 2017). The TEER of the epithelial-endothelial tissue interface was obtained by subtracting the impedance at 100kHz (system background impedance) from that at 100Hz reported as the Organ Chip tissue impedance(Ω). The baseline impedance of empty chips perfused with DPBS was measured to ensure the negligible resistance of the membrane or correction for any background resistance. Throughout the culture, the impedance of siChip was continuously measured every 1-2 days and up to 23 and 12 days for Gut and Liver siChips, respectively.

#### Oxygen measurement and analysis

Oxygen measurements were performed using FireSting®-O_2_ Optical Oxygen and Temperature Meter system (FSO2-C4, Pyroscience sensor technology, Germany) controlled using Pryo Oxygen Logger (V3.317©, Germany). To measure the oxygen levels, siChip was placed on a slide warmer (XH-2002, Premiere®) set at physiological temperature of 37°C while the sensing optical fiber (Pyroscience, Cat. no. SPFIB-BARE) was placed perpendicular to the oxygen sensitive spot on-chip for maximum signal intensity (**Supplementary Fig. S2B**). The oxygen sensor spot was illuminated with red light (610 to 630 nm wavelength) and the luminescence emission was collected at near-infrared (NIR) (760 to 790 nm wavelength) for 10 consecutive seconds (one measurement per second). A fiber optic Pt100 Temperature Probe (Pyroscience, Cat. no. TDIP15) was also used for simultaneous temperature measurement on-chip. The measured signal intensity was converted to dissolved %O_2_ using the Oxygen Calculation Tool (©2019 Pyroscience, Germany). We calibrated the FireSting® system with a 2-point calibration method using 100% and 0% dissolved oxygen calibration solutions following Pyroscience Sensor Technology (Germany) instructions. Briefly, oxygen-sensitive detection spots were immobilized at the flat bottom of two plasma-treated glass containers; one was half-filled with dH_2_O and thoroughly mixed several times to achieve 100% dissolved oxygen (%21 O_2_), and one filled with 0% O_2_ deoxygenated solution made by mixing one OXCAL calibration capsule (Pyroscience) in 50 ml dH_2_O. For Gut and Liver siChip oxygen measurements, representative oxygen tension levels were obtained from the average of 10 consecutive measurements across all of the four sensor locations on-chip.

#### pH measurement and analysis

pH measurements were performed using the FireSting®-PRO Optical PH and Temperature Meter (FRPRO-4, Pyroscience sensor technology, Germany) controlled by Pyroscience Workbench software (V1.2.3.1406, Germany). Similar to oxygen sensing, pH was measured by placing the contactless fiber optic (Pyroscience, Cat. no. SPFIB-BARE) perpendicular to the pH sensor spot, illuminating the spots with the red light through the transparent wall of the reservoir pod and collecting the reflected NIR light over 10 seconds (**Supplementary Fig. S2B**). The measured signal intensity was converted to pH level using the calibration parameters in the Pryo Workbench software. We calibrated the FireSting®-PRO Meter using a 2-point calibration method with pH 2 (PHCAL2, Pyroscience) and pH 10 (PHCAL10, Pyroscience) calibration solutions when pH sensors were mounted on the glass vials.

#### Automated imaging and analysis of metabolic cell activity on-siChip

Metabolic activity of the liver cells was measured continuously using Perceval reporter sensor (lentivirus particle pLV[Exp]-Puro-CMV:T2A:mCherry/3xNLS) (VectorBuilder.com)(Berg et al. 2009) transduced in the Huh-7 cells combined with an in-house developed automated fluorescent microscopy system compatible with the Zoë™ Culture Module. Perceval(ns) ORF and nuclei (mCherry/3xNLS ORF) reporters (Berg et al. 2009) respectively allow for live sensing of the ATP/ADP (adenosine-triphosphate to adenosine-diphosphate) concentration ratio and cell viability. The Perceval sensor offers high-affinity binding site to ATP molecules (prominent peak at 490nm wavelength), but when the ATP levels decrease, ADP molecules (small peak at 405nm wavelength) bind to the same site leading to a binding competition between the two substrates which induces a ratiometric change in the YFP/CFP excitation spectrum that provide a readout for the ATP/ADP ratio. The T2A:mCherry nuclear localization reporter is a live cell marker that uses a peptide produced by living cells that binds to proteins entering the nucleus, which release the fluorescent mCherry reporter. To continuously image the YFP/CFP/mCherry signal in the living chip cultures, we developed a customized microscopy module inside the incubator built on a XYZ translation stage (Zaber Corp) for providing micron positional accuracy along with a CFP/YFP/mCherry multiband filter set (Chroma) that also offers narrow LED excitation bands (405, 505, and 565 nm, Thorlabs). A dock was designed above the microscope to accommodate the Zoë™ with cut through openings that provide optical access to siChips while cultured inside the Zoë™. The microscopy platform was controlled using a Python-based graphical user interface (GUI) developed in-house that allows for the design of the experimental setup, as well as control of the three-axis linear stages, camera components, light settings, automated timelapse imaging, and autofocusing capabilities.

At each timepoint, fluorescent images of the living siChips were acquired automatically at 10 different locations along the length of the apical channel at the three wavelengths (RFP for nuclei and YFP and CFP for ATP/ADP ratio). An automated image processing pipeline was also developed using open-source CellProfiler (cellprofiler.org) software to automatically perform the following analysis on each acquired image: 1) identify nuclei with RFP marker, 2) define a cell as 4 pixels surrounding the edge of each nucleus, 3) measure YFP and CFP signal intensities and the YFP/CFP ratio in each cell, 4) count cell numbers per Image, 5) track changes in YFP and CFP signal intensity over time, and 6) average the YFP/CFP ratio over all cells per image and normalize it to total cell count. ImageJ (Version 1.53) was used to divide the YFP and CFP signal intensities and obtain ATP/ADP images representing metabolic activity changes in the siChip.

### 2.4. Continuous measurement of metabolic functions in response to chemical stimulation and bacteria co-culture on siChip

The baseline levels of TEER, oxygen, pH, and ATP/ADP ratio in siChips were measured up to two days before starting the treatments to ensure the stability of the metabolic activities before any external stimuli were added.

#### Calcium chelation treatment to disrupt cell-cell junctions

Gut and Liver siChips were treated at day 14 and 6 of culture, respectively, with 5 mM ethylene glycol-bis(2-aminoethyl ether)-N, N, N′, N′-tetraacetic acid (EGTA; Sigma, Cat. no. E3889) in dH_2_O that was added to the media of both apical and basal channels and perfused continuously for 30-45 minutes before starting the measurements. The TEER was measured continuously every 20-30 minutes and oxygen was measured every 30 minutes to 1 hour throughout the 2.5-3 hours of the treatment. After 3 hours, the chip media was replaced in both channels with fresh (no EGTA) culture media and the culture was continued for recovery overnight. The impedance and oxygen levels were measured at multiple time points during recovery until normal metabolic functions were resumed on-chip. Bright field and immunofluorescence microscopy imaging was also performed non-invasively throughout the EGTA treatment and recovery time.

#### Oligomycin (OM) treatment

Gut and Liver siChips were treated, at day 17 and 12 of culture, respectively, with 1 µM oligomycin A (Sigma-Aldrich, Cat. no. 75351) in the epithelial and endothelial media perfused continuously in the chip for 3 hours before starting the measurements. Control siChips were treated with media containing 0.5% DMSO (OM drug carrier). After 24 and 5 hours of treatment in Gut and liver siChips, respectively, the media was changed to fresh (non-OM) media to start the tissue recovery. The impedance and oxygen levels were measured continuously during treatment and up to 3 days during the recovery.

Bright field and immunofluorescence (nuclear reporter, ATP/ADP ratio in Huh-7 cells) microscopy imaging was also performed before and during the treatment as well as after the recovery.

#### Lactococcus lactis (L. lactis) bacteria co-culture

*Lactococcus lactis* subsp. *cremoris* MG1363 (*L. lactis*) (ATCC, Cat. no.19257) was inoculated in the apical epithelium channel of the Gut Chip at day 13 of culture. To remove any antibiotic or buffering capacity from the media, the chips were perfused with a customized low buffer Hanks Buffer Salt Solution (HBSS, pH 5.1) in the apical channel and with antibiotic-free media in the basal channel 24 hours prior to bacterial inoculation. The customized HBSS solution allows modulation of the pH during the co-culture time. The chips were inoculated with *L. lactis* (10^3^ CFU/ml) at 37°C for 1-2 hours under static conditions before resuming continuous flow on the chip in the Zoë™ Culture Module. *L. lactis* co-culture on Gut Chip was maintained for 4 days and the pH level of the apical and basal endothelium channels were continuously measured every 24 hours.

#### Immunofluorescence staining and imaging

At the endpoint of experiments, siChips were removed from the mPods, washed two times with DPBS, fixed with 4% paraformaldehyde for 15-20 minutes at room temperature, and incubated overnight in 1:100 anti-H2AX antibody (Novus Bio NB100-2280) at 4°C in 1% BSA in PBS and 0.01% Triton-X 100. The chips were washed with PBS containing 0.01% Triton-X 100 twice before incubation in the secondary antibody (1:1000 anti-rabbit 555) and 1:10,000 DAPI, and imaging on a Zeiss LSM 710 with a 32x water objective.

### 2.5. Statistical analysis

All results are presented from at least two independent experiments and data points are shown as mean ± standard error of the mean (s.e.m.) from multiple chips. Statistical analysis was performed using GraphPad Prism version 8.0 (GraphPad Software Inc., San Diego, CA) and the statistical significances between groups were tested using two-tailed Student’s t-test, unpaired multiple t-test or One-way ANOVA with Tukey correction for multiple hypothesis testing.

## 3. Results and Discussion

### 3.1. Fabrication of the siChips with TEER, oxygen, and pH sensing capabilities

The siChip was designed to enable three main capabilities: i) recapitulation of the tissue physiological structure and microenvironment, ii) non-invasive and continuous sensing of multiple metabolic functions, and iii) compatibility with a commercially available tissue culture module for automated and multiplexed Organ Chip culture. The siChip was fabricated using a layer-by-layer assembly approach that incorporated flexible PDMS polymer for making apical and basal channels as well as a porous membrane to allow implementation of biomechanical stimulation (e.g., peristalsis-like motions) on-chip. Rigid PC material was also used and integrated with TEER and oxygen sensors to minimize gas permeability and measure epithelial barrier function and oxygen levels directly inside the chip microenvironment (**Fig. 1A)**. The integrated sensors that detect free oxygen molecules present in the pericellular microenvironment of the tissue were placed at four different locations in the siChip (two in apical and two in basal channels) to fairly represent tissue oxygen tension levels across the whole chip. These oxygen sensing spots are easily and non-invasively accessible through the optically clear PC layer on the top and bottom sides of the siChip allowing continuous, real-time, oxygen measurements in living chip cultures throughout the entire experimental timeline (**Fig. 1B**, **Supplementary Fig. S1C**).

**Figure 1.**
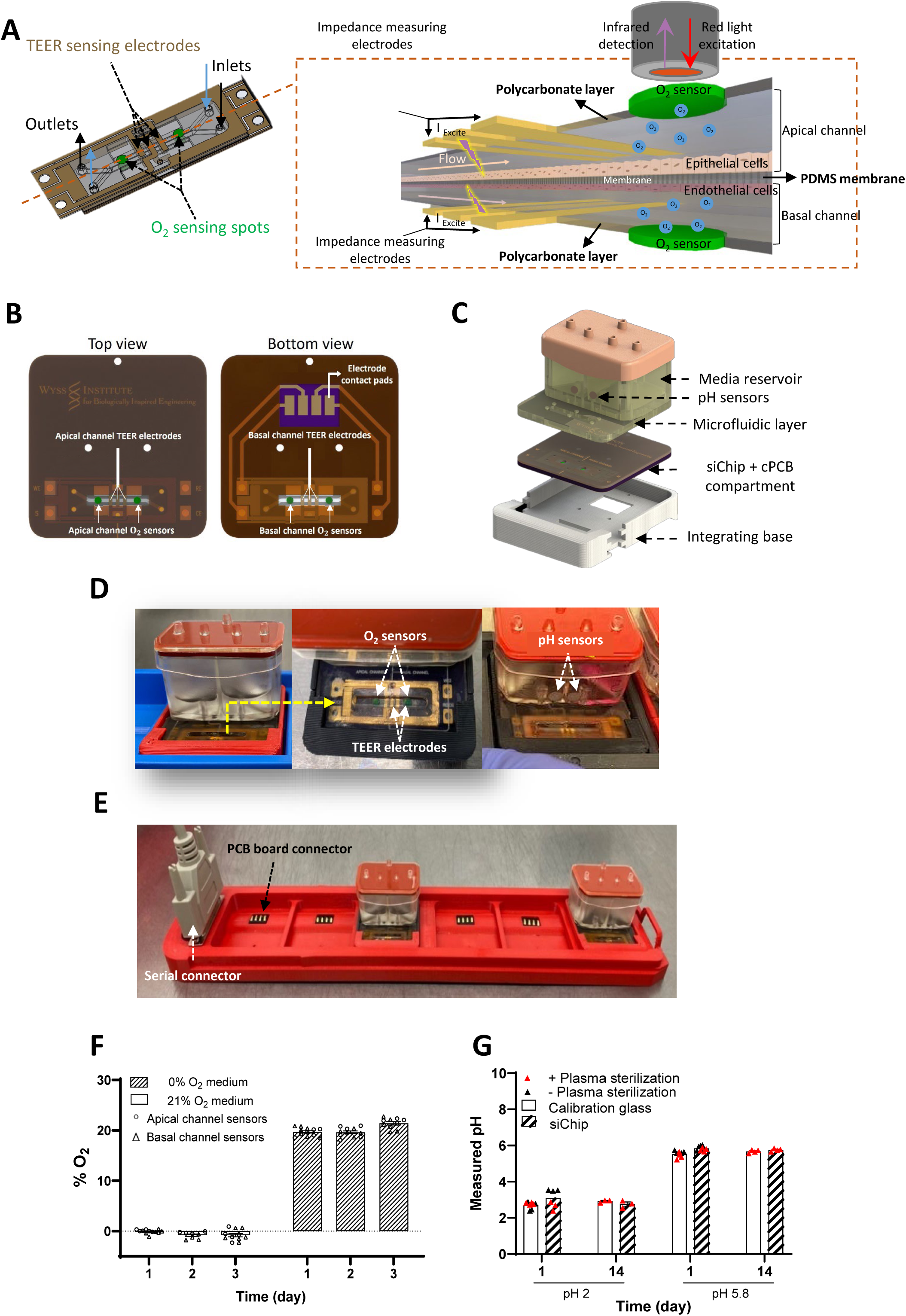
Design, fabrication, and validation of measurements in sensor integrated Organ Chip (siChip). **A.** Schematic view of the dual channel chip with integrated TEER and oxygen sensors (left) and magnified cross section view of the chip (right). **B.** Schematic of the top and bottom views of the assembled siChip on customized PCB (cPCB). **C.** Assembly of the siChip on cPCB with Pod^TM^ media reservoir to make a modified pod (mPod). **D.** Photographs of the final siChip device with integrated TEER, oxygen, and pH sensors for automatic flow perfusion and culture in Zoe™ Culture Module. **E.** Photograph of the modified Tray (mTray) with pin connectors for continuous and remote TEER measurements. **F.** Validation of oxygen sensors measurements in siChip with 0% and 21% O_2_ calibrating solutions. **G.** Validation of pH sensors measurement before and after siChip sterilization as compared to standard calibration glass using pH 2 and pH 5.8 solutions. n=6 (**F**) and 4 (**G**) individual siChips per group.

The integrated TEER sensors were used to measure barrier function in the siChip remotely and continuously. A customized PCB (cPCB) with compatible configuration to the outline of Pod^TM^ organ chip holder (Emulate Inc.) was developed that allows for both direct access to the TEER electrodes and seamless integration of the siChip into the Pod^TM^ (**Fig. 1B**). We designed and 3D printed a base compartment that facilitates this by attaching the siChip-on-cPCB to the modified Pod^TM^ (mPod) (**Fig. 1C, Supplementary Fig. S1F**). In addition, we integrated pH sensors in close vicinity of the outlets of the epithelial and endothelial channels to measure pH levels within the effluents of each channel. The designs of the mPCB and mPod enabled seamless integration of the siChip device within the commercially available, automated Zoë^TM^ Organ Chip culture module. The design also allowed for non-invasive access to all the integrated sensors (e.g., TEER, oxygen, pH) through an opening in the mPod that overlaid the channels of the chip, which permitted continuous measurements and frequent optical imaging and metabolic monitoring of live chip cultures (**Fig. 1D, Supplementary Fig. S2,B**). We also developed a modified Tray (mTray) that provides direct and simultaneous access to the mPCB of 6 siChips in parallel and hence, remote measurement of TEER while the siChips are being cultured inside the Zoë^TM^ culture instrument. Each mTray consisted of an array of built-in spring-loaded electrical contacts in the base of each slot for direct contact with the cPCB in each siChip. Using a serial connector, the electrical signals sensed from all siChips are transferred out via a serial cable enabling remote data collection from outside the incubator (**Fig. 1E, Supplementary Fig. S2A**). We used a combination of in-house customized designs and fabrication techniques with a commercially available system to simultaneously maximize the non-invasive and functional sensing capabilities, and user-friendly operation of the siChip device as well as scale-up manufacturing potential, biocompatibility and clinical relevance.

We next tested the stability of the integrated oxygen and pH sensors in siChips without cells for accurate, reproducible, and long-term stable measurements of tissue metabolic activity on-chip. The siChips were perfused with solutions of known oxygen or pH levels using the Zoë^TM^ Culture Module operated inside the incubator (37°C, 5%CO_2_) for 3-14 days while measuring oxygen and pH levels. Continuous perfusion of 0% and 100% (21% O_2_) air saturated solutions in the siChips and on-chip measurements confirmed the accuracy and stability of the oxygen tension signals as well as the sensors’ signal consistency in the apical and basal channels (**Fig. 1F**). The long-term stability and accuracy of the pH sensors under tissue culture conditions over time were also confirmed using pH 2 and 5.5 solutions perfused through the siChips for 14 days. Results obtained using the on-chip pH sensors and in standard calibration glass containers were comparable. Additionally, oxygen plasma treatment (used for siChip sterilization before cell culture) did not affect the sensor’s functionality (**Fig. 1G**). These results collectively indicate that the bespoke siChip and integrated analytical sensors maintain their integrity, function, and measurement accuracy during long time experiments under physiologically relevant tissue culture conditions.

### 3.2. Non-invasive monitoring of metabolic functions in Gut and Liver siChips

After fabrication of the siChips with TEER, oxygen, and pH sensing capabilities, we validated their biocompatibility with living cells and functionality by creating Gut and Liver Chips using the siChip devices and monitoring their viability and metabolic activity. In the Gut siChip, human Caco-2 intestinal epithelial cells and human umbilical vein endothelial cells (HUVECs) were cultured on the apical and basal surfaces of the ECM-coated membrane, respectively, to create a physiologically relevant epithelial-endothelial tissue interface, which recapitulates many features of the human small intestine, including intestinal villi formation, drug metabolizing activity, and mucus production, as described previously (Jalili-Firoozinezhad et al. 2019; Kim et al. 2012). As observed in past Gut Chip studies, the epithelial cells formed a uniform monolayer at day 1, developed into a dense villus epithelium by day 10, and accumulated a thick mucus layer by day 23 that obscured the epithelium below (**Supplementary Fig. S3A**).

A Liver siChip was similarly developed by seeding Huh7 liver epithelial cells that are well characterized (Kawamoto et al. 2020) and liver sinusoidal endothelial cells (LSECs) on the apical and basal sides of the chip porous membrane, respectively. Phase contrast images of the chips showed formation of a uniform cell monolayer at day 1 that formed a dense tissue by day 4, 8, and 12 on siChip (**Supplementary Fig. S3B**), similar to previously reported established Liver Chip that used Huh7, HepG2 or hiPSC-derived hepatocytes (Farooqi et al. 2021; Kulsharova et al. 2021; Lee-Montiel et al. 2020; Menon et al. 2014).

#### Continuous measurement of barrier function on siChips

Barrier function plays an important role in epithelial tissues in health and disease and thus, we used our integrated TEER electrodes as a label-free, noninvasive way to analyze barrier properties (impedance) of the Gut and Liver siChips every 1 to 2 days over more than 3 weeks in culture while under continuous flow on-chip. In the Gut siChips, we detected an electrically resistant intestinal barrier with an impedance of > 2,000Ω at day 1 of culture, which remained relatively constant while the chips were perfused with the growth medium during the first 10 days of culture; however, the impedance gradually increased (up to >1.5 fold) when the serum level was dropped in the perfused medium to induce cell differentiation (**Fig. 2A**). The higher levels of electrical impedance during the differentiation period compared to expansion period are consistent with past studies that showed an increase in epithelial tight junctions during the differentiation process (Varadarajan et al. 2019).

**Figure 2.**
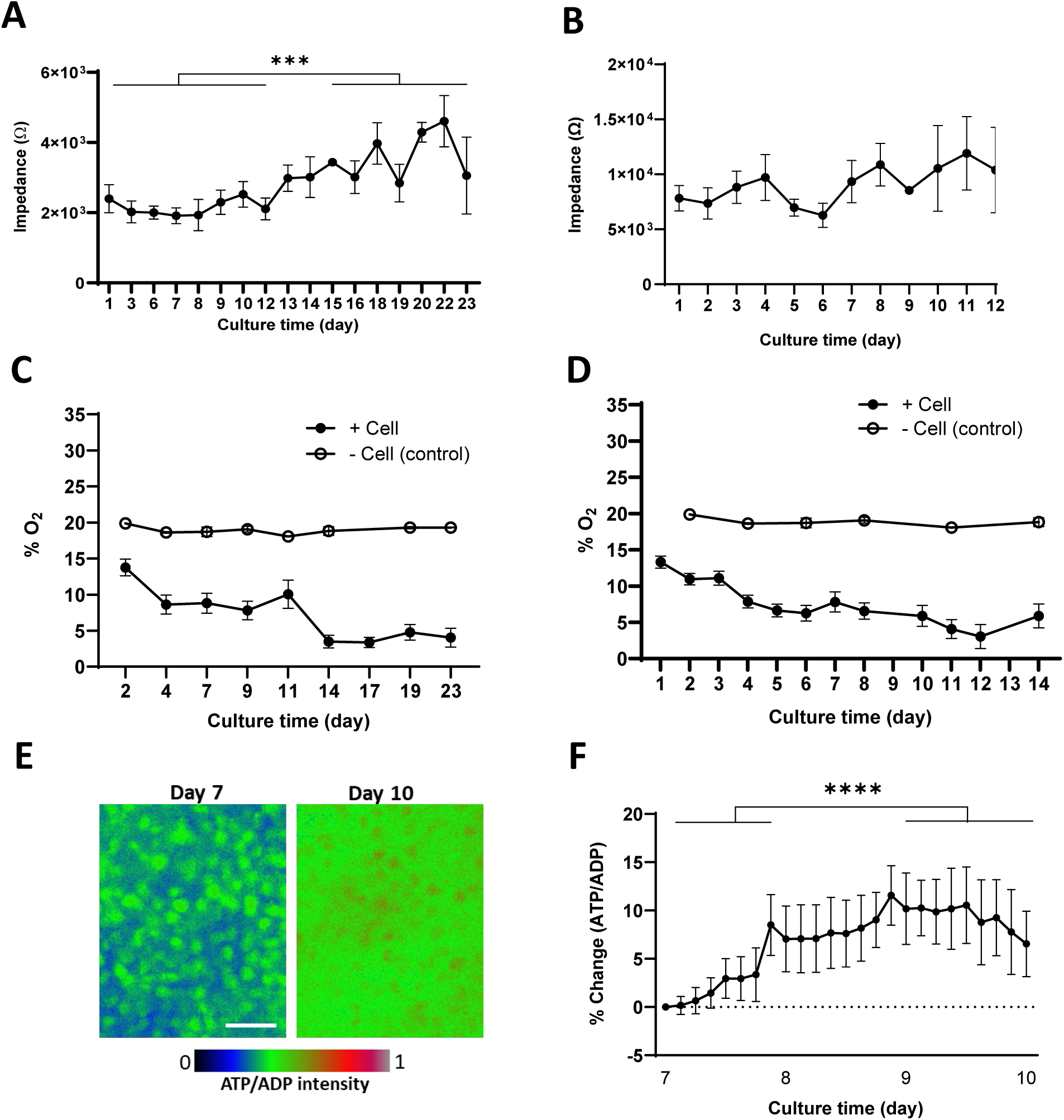
Non-invasive monitoring of metabolic functions in Gut and Liver siChips. Continuous measurement of **A, B.** impedance, and **C, D.** oxygen tension in Gut (**A,C**) and Liver (**B,D**) siChips throughout the culture time. **E.** Fluorescence ATP/ADP signal microscopic images at day 7 and 10 of culture in Liver siChip. Scale bar 100 μm. **F.** Continuous measurement of percentage change in ATP/ADP fluorescence signal in Liver siChip over 3 days of culture. Data represent the mean ± s.e.m.; n=4-9 (**A**), 5-9 (**B**), 5-13 (**C**), 15-25 (**D**), and 5 (**E**) experimental chip replicate. *\*\*\* P< 0.001, **** P< 0.0001*.

The continuous measurement of TEER in Liver siChip similarly showed the establishment of an electrically resistance (>7,800Ω) tissue barrier at day 1 of culture which increased up to >1.2 fold by day 4 and remained relatively stable during the rest of the 12 day experiment when a dense liver epithelium was established on-chip (**Fig. 2B**, **Supplementary Fig. S3B**).

Electrical impedance encompasses both resistive and capacitive properties of the tissue barrier. At a frequency range of 100Hz to 10kHz, the current cannot pass through the tissue layer and the measured impedance reflects the combined effect of cell-cell junctions in the epithelium and endothelium as well as the stored current in the cells, which represents the tissues’ electrical capacitance (Morgan et al. 2019; Srinivasan et al. 2015). At the 100Hz frequency used in this study, the measured impedance represents the tissue electrical resistance (Henry et al. 2017).

#### Continuous measurement of oxygen levels and ATP/ADP activity on siChips

Respiration is one of the key indicators of the cell and tissue metabolic activity and quantification of dissolved oxygen levels in culture medium can improve our understanding of tissue cellular functions in normal and disease states (Johannsen and Ravussin 2009; McClelland 2011). In addition, having a continuous measure of oxygen levels would enhance consistency and reproducibility in culture studies (Al-Ani et al. 2018). We leveraged the oxygen sensors that we integrated into the siChips (**Fig. 1A**) to measure the dissolved oxygen levels in the medium flowing through the epithelial channels of the Gut and Liver siChips every 1-3 days throughout the culture to monitor the epithelial cell respiration activity. In the Gut siChips, continuous oxygen measurements under culture conditions revealed that the oxygen level drops to <15% at day 2 and <10% at day 4 of culture based on cellular utilization of oxygen (Grant et al. 2022), after which it remains stable until day 11 corresponding to the end of the expansion period in culture. Changing the expansion medium to the low serum differentiation medium further reduced the oxygen tension to microaerophilic levels (<5%) by day 14, and this remained stable for the rest of the culture time (**Fig. 2C**). This finding is consistent with a higher rate of aerobic respiration (>80% drop in oxygen tension) in the dense and differentiated epithelium compared to proliferating epithelial cells (**Supplementary Fig. S3C**).

Continuous measurements of oxygen level in the Liver siChip revealed that the tissue reduces the oxygen levels in the medium to <15% as early as day 1 in the culture, and this continues to drop reaching <5% by the end of the cell expansion period at day 5. As the Liver siChip continues to grow during the differentiation time, the oxygen level stabilizes at ∼5-7% with >80% drop in oxygen level compared to day 1 (**Fig. 2D, Supplementary Fig. S3D**).

We also continuously measured the metabolic activity and viability of the epithelial cells expressing reporters for ATP/ADP and T2A:mcherry NLS (nuclear localization signal) in the Liver siChips by measuring the fluorescent intensity of the ATP/ADP ratio and cell nuclear signals, respectively. Continuous fluorescent imaging of siChips every 30-60 minutes showed an increase in the ATP/ADP ratio at day 10 compared to day 7 of culture and revealed a 6.5% increase in liver cell metabolic activity over 3 days of culture during differentiation (**Fig. 2E,F**). Continuous monitoring of the RFP nuclei signals also showed that the Huh7 cells remain viable on-siChips throughout the 12 days of culture (**Supplementary Fig. S3E**). The Huh7 liver epithelial cell line was originally isolated from a liver carcinoma and it is known to have low enzyme and metabolic activity compared to primary human hepatocytes (Bulutoglu et al. 2020), but it can become more metabolically active as it can continue growing past confluence over long periods of culture (up to 4 week) (Sivertsson et al. 2010). Transfecting the cells with fluorescent reporters provided additional continuous monitoring capabilities in the siChips that enabled both direct and quantitative measurement of cell metabolic activity and viability, as well as automated image focusing and acquisition that excludes dead cells from analysis.

In the body, intestine and liver organs maintain physiological homeostasis under microaerophilic oxygen tension; 0.4-10% O_2_ in the intestinal lumen (He et al. 1999) and 5-8% in the liver (Carreau et al. 2011; Guo et al. 2017; Trepiana et al. 2017). Recapitulating similar microaerophilic oxygen tension in the Organ Chips has been challenging in chips made of PDMS (Jalili-Firoozinezhad et al. 2019; Shin et al. 2019) due to the constant diffusion of the ambient air into the tissue channels that dominates any micro-modulation of the oxygen level by the cell’s aerobic respiration. Here, we used thermoplastic PC polymer with limited oxygen permeability (Byrne et al. 2014; Mehta et al. 2009) in the top and bottom walls of the siChip to create a semi-isolated microenvironment for cells so that they can physiologically modulate the oxygen levels to microaerophilic levels on-chip through aerobic respiration similar to that *in vivo* (Guo et al. 2017; He et al. 1999). This design further allowed us to model the effect of temperature on aerobic respiration in the intestinal epithelium, which revealed a higher oxygen consumption rate (up to 73%) in the Gut siChip at physiological temperature (37°C) compared to room temperature (25 °C) (**Supplementary Fig. S3F**). The oxygen uptake in the ileus has been similarly reported to correlate with the lumen temperature consistent with the change in the metabolic rate (Kvietys et al. 1985). A temperature drop from 32 to 22°C was reported to reduce oxygen consumption in the kidney and intestinal tissue by 75% (Hendriks et al. 2019; Kvietys et al. 1985), which is also similar to our observation in the Gut siChip further confirming the robustness of our system for modeling and sensitively monitoring tissue physiological responses to microenvironmental changes. The less permeable PC walls of the siChip also prevent absorption and adsorption of small, hydrophobic molecules (i.e. hormones and drugs) to the chip’s bulk material (Auner et al. 2019), which can be a challenge in using PDMS chips, particularly studies with highly lipophilic chemicals or drugs (Toepke and Beebe 2006). Using flexible PDMS material in the main body of the siChip still allows for the application of cyclic biomechanical forces (i.e. peristalsis) when essential for tissue differentiation (Huh et al. 2010; Kim et al. 2012), which is an advantage compared to fully thermoplastic devices that cannot accommodate this functionality (Azizgolshani et al. 2021; Bussooa et al. 2022). Overall, our siChip design eliminates current technical challenges in creating physiological oxygen levels on-chip including anoxic medium perfusion in the channels or chip isolation inside an anaerobic chamber (Jalili-Firoozinezhad et al. 2019; Shin et al. 2019), while providing a confined microenvironment for cells to naturally establish relevant tissue/organ physioxia *in vitro*. These advancements expand the use of Organ Chip *in vitro* models in diverse and potentially higher throughput preclinical studies.

### 3.3. Continuous monitoring of responses to chemical and bacterial stimuli

#### Dynamic monitoring of Gut siChip responses to environmental stimuli

To evaluate the sensitivity of the siChip system for continuous monitoring of functional and metabolic changes in the Organ Chips, we treated Gut siChips with chemicals and/or bacteria that are known to modulate epithelial tissue barrier, cell respiration, and pH. EGTA is a high affinity calcium chelator that can reversibly disrupt epithelial tight junctions by reducing free calcium concentration in the culture medium (Rothen-Rutishauser et al. 2002; Samak et al. 2011). We exposed the Gut siChips to 5 mM EGTA for 3 hours at day 17 of culture when the intestinal tissue had attained phenotypic and metabolic functional stability, and continuously measured TEER and oxygen levels during the treatment period and for up to 24 hours after the EGTA was removed from the medium. Continuous on-chip TEER measurements showed >80% decrease in the impedance (from > 2,000Ω to < 800Ω) within 2 hours after exposure to EGTA, which recovered to the pre-treatment level (>2,400Ω) that also was similar to that measured in control Gut siChips within 12 hours after removing the EGTA (**Fig. 3A**). Phase contrast microscopic imaging of the siChips revealed disruption of the opaque mucosal layer and retraction of the epithelium resulting in the appearance of large, rounded structures 3 hours after exposure to the EGTA. Normal epithelial tissue morphology was fully recovered by 4 days after the EGTA was removed (**Supplementary Fig. S4A**).

**Figure 3.**
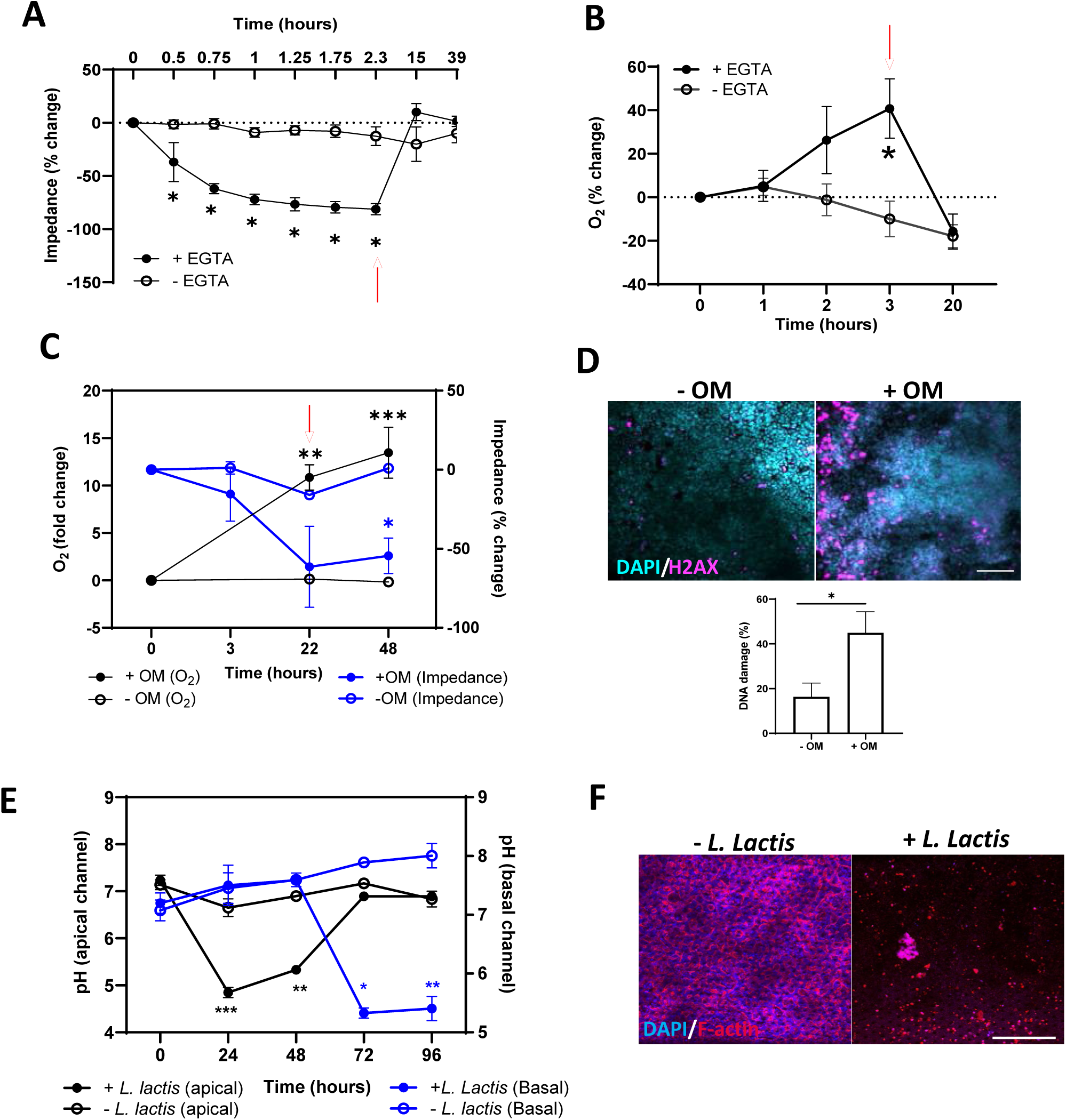
Continuous monitoring of Gut siChip functional and metabolic responses to chemical and bacterial stimuli. **A.** Percentage change of impedance in Gut siChip with time during EGTA treatment and after it was removed. **B.** Continuous measurement of percentage change in oxygen level during EGTA treatment and post recovery. **C.** Change in the Gut siChip oxygen level (black line) and impedance (blue line) in response to oligomycin (+OM) or its vehicle (-OM) exposure and after they were removed. **D.** Immunofluorescence images (top) of siChip intestinal epithelium 48 hours after treatment with OM compared to drug carrier stained with DAPI (nuclei, blue) and H2AX (DNA damage, magenta). Image-based quantitative analysis of percentage change in epithelial cells DNA damage with vehicle and OM treatment (bottom). Scale bar 100 μm. **E.** Measurements of pH changes in outflows of epithelium (black line) and endothelium (blue line) lumens in response to *Lactococcus Lactis* (*L. lactis*) bacteria co-culture. **F.** Immunofluorescent microscopy images of the gut epithelium stained with DAPI (nuclei, blue) and F-actin (magenta) markers 96 hours after inoculation with *L. lactis* bacteria. Scale bar 100 μm. Data represent the mean ± s.e.m.; n=5 (**A**), 3 (**B**), 3-6 (**C**), and 3-6 (**E**) experimental chip replicate. *\* P< 0.05, ** P< 0.01, *** P< 0.001.* Presented percentage and fold change values are normalized to pre-treatment levels. Red arrows indicate the end of EGTA or OM treatment in **A,B,** and **C.**

Calcium chelating agents are known to modulate mitochondrial respiration in cells (Masola and Evered 1984). Simultaneous measurement of oxygen levels on-chip revealed no change in the tissue respiration rate during the first hour of the treatment despite significant reduction (>70%) in the barrier function. However, after 3 hours, oxygen levels had significantly increased up to 40%, representing a significant drop in the tissue aerobic respiration that reversed back to control, non-treated chip levels 17 hours after removing the EGTA (**Fig. 3B**). The observed effect of EGTA on intestinal epithelial barrier resistance is also similar to that reported in previous studies (Azizgolshani et al. 2021; Henry et al. 2017). However, the multiplexing of the integrated TEER and oxygen sensors revealed simultaneous dynamic effects of EGTA on both barrier function and aerobic respiration in the intestinal tissue *in vitro*, which has not been reported before.

Oligomycin (OM) is a well-characterized antibiotic that binds to subunit of the mitochondrial ATP synthase and can significantly affect mitochondrial-dependent ATP production and cell respiration (Hao et al. 2010; Lee and O’Brien 2010; Ruas et al. 2016). At day 17 of culture, OM (1 μM) was perfused through the Gut siChip for 22 hours and then removed while carrying out continuous on-chip monitoring of oxygen levels and TEER. Continuous oxygen measurements showed a significant (10-fold) increase in oxygen level after 22 hour treatment with OM, which further increased to >12-fold even after the OM was removed from medium, suggesting that some of the damage to epithelial cell respiration function was non-reversable (**Fig. 3C**).

Continuous TEER measurements also demonstrated an adverse effect of OM on barrier function as indicated by a 25%-60% decrease in barrier impedance during exposure to OM that did not return to normal levels after termination of the OM treatment (**Fig. 3C**).

Immunofluorescent microscopic imaging of the chip 22 hours after treatment and 24 hours after the recovery, respectively, showed an increase in the number of H2AX positive cells (>18%), indicating the presence of significant DNA damage in the chip epithelium, compared to untreated control chips (**Fig. 3D**). These adverse effects in DNA damage and oxygen respiration observed on-chip are consistent with previous studies that reported the detrimental effects of OM exposure in Caco-2 intestinal epithelial cells and HUVEC (JanssenDuijghuijsen et al. 2017; Patella et al. 2015).

pH homeostasis is also highly relevant for human tissue and organ physiology, as well as host interactions with bacterial and viral microorganisms (Espinosa 2002; Hackett et al. 2016; Perdikis et al. 1998). Changes in pH have been observed in different diseases (Aoi and Marunaka 2014; Lykke et al. 2021; Nakano et al. 2015) and thus, continuous quantitative measurement of pH *in vitro* can be a powerful tool to better understand dynamic tissue responses to different stimuli and insults. Because microbial interactions with host epithelium contribute significantly to control of tissue pH under both physiological and pathophysiological conditions (Jalili-Firoozinezhad et al. 2019; Perdikis et al. 1998; Plesniarski et al. 2021), we used our Gut siChip with integrated pH sensors to continuously monitor pH levels when populated with living bacteria. On culture day 17, the chip was inoculated with *Lactococcus lactis* (*L. lactis),* a non-pathogenic, gram-positive bacterium that uses lactose or pyruvate for ATP formation and subsequent lactic acid production, which is known to dramatically decrease pH in the tissue microenvironment (Hugenholtz et al. 1993). Continuous pH measurements on-chip indicated that co-culture with the bacteria for 1 to 2 days resulted in a significant reduction in medium pH (from 7.2 to 4.5) in effluents of the epithelial channel, while the pH of the control chips without bacteria was maintained at a constat level (∼pH 7) (**Fig. 3E**). In contrast, the medium pH of the endothelial effluents remained unchanged during this same period, indicating that intestinal tissue barrier that prevented bacteria from entering the endothelium lumen remained intact. After 2 days, the pH of the endothelial medium dropped significantly to < pH 5, which is consistent with disruption of the tissue barrier resulting in invasion of *L. lactis* bacteria into the underlying endothelium. This was associated with a change in the color of phenol red containing endothelium medium from red (∼ pH 7.4) to yellow (acidic pH) with observable cloudiness representing overgrowth of bacteria in the medium (**Supplementary Fig. S4B**).

Interestingly, the pH drop in the endothelial channel at these later times was accompanied by a simultaneous increase in the pH of the epithelium lumen (**Fig. 3E**). This may partially be explained by the overgrowth of bacteria in the epithelial channel resulting in a major depletion of nutrients and epithelial cell death and detachment at later times, which is consistent with results of immunofluorescence microscopic imaging of the epithelial layer (**Fig. 3F**). Once the barrier disrupted, the bacteria may move to the endothelial channel where higher nutrient concentrations are present leading to reduced numbers of lactic acid producing bacteria in the epithelial channel, and hence increased pH.

#### Dynamic monitoring of Liver siChip responses to environmental stimuli

Similar to the Gut siChip, we exposed differentiated Liver siChips to EGTA (5 mM) for 3 hours and continuously monitored oxygen tension and barrier impedance on-chip throughout the treatment and recovery periods. Oxygen levels in the Liver siChip significantly increased and reached up to 80% higher values than before the treatment within 3 hours of exposure indicating that aerobic respiration progressively decreased in the liver tissue with EGTA treatment, similar to what we observed in the Gut siChip. Moreover, removal of the EGTA restored normal tissue respiration and reduced the oxygen level back to control levels within 21 hours (**Fig. 4A**).

**Figure 4.**
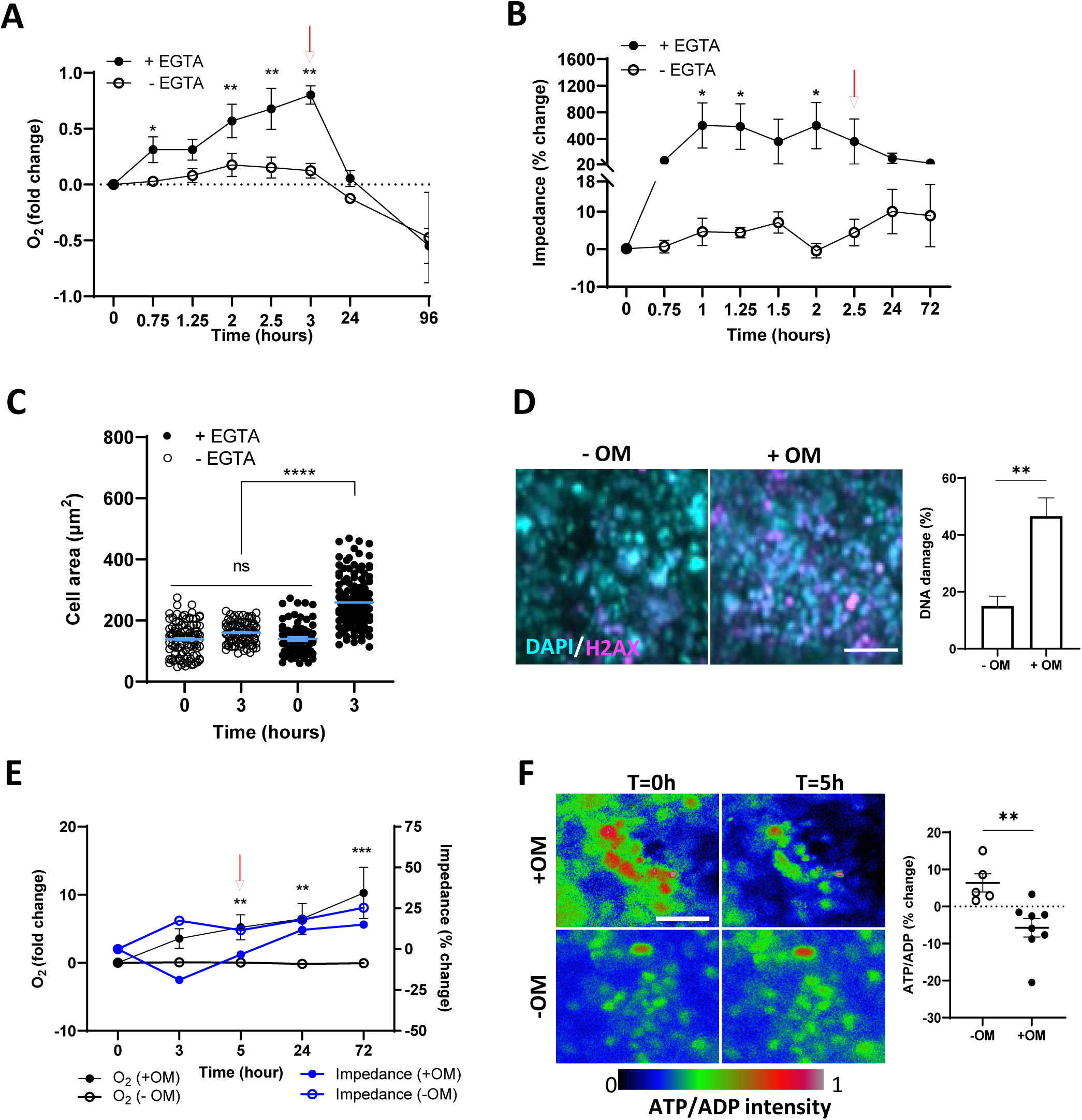
Continuous monitoring of Liver siChip responses to chemical stimuli. Dynamic fold change in oxygen level (**A**) and percentage change in impedance (**B**) of Liver siChip throughout exposure to EGTA and after its removal. **C.** Quantitative analysis of epithelial cell size before and after 3 hours of EGTA treatment compared to control chips with no EGTA. **D.** Immunofluorescent microscopic images (left) and quantitative analysis (right) of the DNA damage (H2AX^+^ cells, magenta) in epithelium after exposure to OM compared to drug carrier. Nuclei stained with DAPI (blue). Scale bar 100 μm. **E.** Dynamic changes in oxygen level (black line) and barrier impedance (blue line) after OM treatment and its removal compared to drug vehicle treated (-OM) siChips. Presented percentage and fold change values are normalized to pre-treatment levels. **F.** Representative images of ATP/ADP fluorescent signal intensity (left) and image-based quantitative analysis (right) before and after 5 hours of exposure to OM (+OM) or drug vehicle (-OM). Scale bar 100 μm. Data represent the mean ± s.e.m.; n=5 (**A**), 5 (**B**), 10 (**C**), 6-15 (**E**) and 6 (**F**) experimental chip replicate *\* P< 0.05, ** P< 0.01, *** P< 0.001, **** P< 0.0001.* Red arrow indicates the end of EGTA or OM treatment in **A, B**, and **E.**

Interestingly, however continuous impedance measurements revealed an unexpected increase in barrier resistance (up to 6-fold) one hour after the EGTA exposure that remained stable throughout the 2.5 hours of the treatment and gradually dropped back to levels comparable to control chips after removing the EGTA at 24 and 72 hours (**Fig. 4B**). This was accompanied by a greater than 1.5-fold increase in Huh7 epithelial cell size (projected area) compared to control untreated chips, as detected by live cell imaging (**Fig. 4C and Supplementary Fig. S4C**). The effects on TEER may be in part explained by an increase in cell proliferation as well as a decrease in cell motility mediated through store-operated calcium entry (SOCE) (Selli et al. 2015), which also has been shown to correlate with higher TEER levels in Huh7 cells (Morgan et al. 2019; Yang et al. 2013). Increased cell spreading enhances growth of many cells (Chen et al. SCIENCE 1997), including Huh 7 cells, while the opposite is true for migratory Huh7 cells (Berger et al. 2015). SOCE inhibition also has been associated with lower mitochondrial respiration (González-Sánchez et al. 2017), which is also consistent with the higher oxygen tension we observed during EGTA treatment in the Liver siChip (**Fig. 4A**).

Additionally, we lowered the serum concentration of the medium 2 days before exposure to EGTA to induce tissue differentiation, and the proliferative capacity of Huh7 cells in the presence of EGTA has been previously shown to be higher when they were pre-conditioned with serum-starvation compared to medium with high serum (Modica et al. 2019).

Similar to the Gut siChip, when we treated the Liver siChip with OM, we observed an increase in the number of DNA-damaged cells after 6 hours and greater DNA damage 3 days after OM removal compared to control chips treated with the diluent (0.5% DMSO) alone (**Fig. 4D**). We also detected a significant (5-fold) increase in oxygen levels within 5 hours after OM addition, which increased up to 10-fold even after OM removal, again indicating a reduced aerobic respiration capacity that was irreversible. OM treatment was also associated with a decrease in barrier impedance 3 hours post-treatment but it recovered to baseline pre-treatment levels 3 days after OM removal (**Fig. 4E)**. Using fluorescent reporters indicating ATP/ADP metabolic activity and cell viability, we observed >12% drop in the ATP/ADP signal ratio at 5 hours and >14% drop in cell viability at 3 hours after treatment compared to control chips (**Fig. 4F, Supplementary Fig. S4D**). These findings are consistent with past studies that showed OM produces a similar decrease in oxygen consumption in Huh7 cells (Theurey et al. 2016; Turcios et al. 2019; Zeidler et al. 2017) as well as in endothelial cells (Patella et al. 2015). In the Huh7 cells, OM also has been shown to reduce cell proliferation and increase cell death (Nwosu et al. 2018). This is consistent with our finding that OM treatment resulted in decreases in tissue barrier function and anaerobic respiration, as well as an increase in the DNA damage and cell viability in the Liver siChips. Moreover, this decrease in proliferation and oxygen consumption was mirrored by a significant drop in metabolic activity evidenced by a decreased ATP/ADP ratio in OM-treated chips. Interestingly, the Liver siChip impedance recovered after 5 hours of exposure while the oxygen tension continued to increase, which in combination may suggest that inhibition of mitochondrial respiration induces compensatory activities in cells to maintain monolayer and barrier integrity; however, confirming of this hypothesis will require future studies.

Previous reports of microfluidic organ chip devices with integrated electrical, electrochemical, and optical sensors (An et al. 2019; Azizgolshani et al. 2021; Fuchs et al. 2021; Henry et al. 2017; Jalili-Firoozinezhad et al. 2019; Li et al. 2017; Lind et al. 2017; Maoz et al. 2017; Müller et al. 2021; Odijk et al. 2015; Riahi et al. 2016; van der Helm et al. 2019; Zhang et al. 2017) have shown their potential value for different tissue monitoring applications. However, these past studies offered limited on-chip analytical sensing capabilities, had complicated designs that impeded multiplexing and automation, or lacked integration of biomechanical cues such as flow for physiological modeling of 3D tissue structure and function. Our siChip system with integrated, multifunctional, analytical sensing capabilities tackled these challenges by incorporating electrical, chemical, cellular, and optical sensing and monitoring capabilities for continuous non-invasive measurement of TEER, oxygen tension, pH, cell phenotype and metabolic activities in one device while enabling simultaneous live optical imaging of the chip through the entire duration of the culture. In addition, it allows for easy and reproducible fabrication, scale-up, and multiplexing, while being compatible with a commercially available automated Organ Chip culture instrument. siChip technology can enable establishment and maintenance of tissue microenvironments with physiological oxygen tension levels, and facilitate more comprehensive studies to improve our understanding of human tissue and organ physiology and disease states.

## 4. Conclusion

Here we presented a multi-sensor integrated Organ Chip (siChip) system that provides a robust platform for continuous and simultaneous measurement of different metabolic functions *in vitro*, as demonstrated in human Gut and Liver siChip over extended culture times. We showed that the siChip design made of both PC and PDMS materials allows us to recapitulate *in vivo-*like morphological, oxygen tension, metabolic, and functional human tissue responses to chemical and bacterial stimuli, while allowing near-continuous analytical measurements of these responses at high temporal resolution. The straightforward and scalable fabrication of siChips, their operation both as standalone system as well as modules that can be cultured using commercially available automated culture instruments, and their inline remote data collecting interfaces facilitate their applicability for drug and therapeutic development and testing with clinically relevant reproducibility and reliability.

## Credit authorship contribution statement

**Zohreh Izadifar:** Conceptualization, Methodology, Investigation, Validation, Resources, Formal analysis, Writing – Original Draft & Editing, Visualization, **Berenice Charrez:** Investigation, Resources, Formal analysis, Writing – Original Draft & Editing, Visualization, **Micaela Almeida:** Methodology, Investigation, Resources, Visualization, **Stijn Robben:** Investigation, Software, **Kanoelani Pilobello:** Investigation, Resources, **Janet van der Graaf-Mas:** Investigation, Software, **Max Benz:** Methodology, Resources**, Susan L. Marquez, Thomas Ferrante:** Resources, **Kostyantyn Shcherbina:** Software, **Russell Gould:** Data Curation, **Nina T. LoGrande:** Resources, **Adama M. Sesay:** Conceptualization, Methodology, Writing – Review and Editing, Supervision, Project administration; **Donald E. Ingber:** Conceptualization, Funding acquisition, Writing – Review and Editing, Supervision.

## Acknowledgment

We acknowledge research funding from the Bill and Melinda Gates Foundation, Seattle, WA (INV-035977 to D.E.I.) and DARPA (W911NF-19-2-0027 to D.E.I.), Canada’s Natural Science and Engineering Research Council (NSERC Postdoctoral Fellowship to Z.I.), and support from the Wyss Institute for Biologically Inspired Engineering at Harvard University.

## SUPPLEMENTARY FIGURE LEGENDS

**Supplementary Figure S1.**
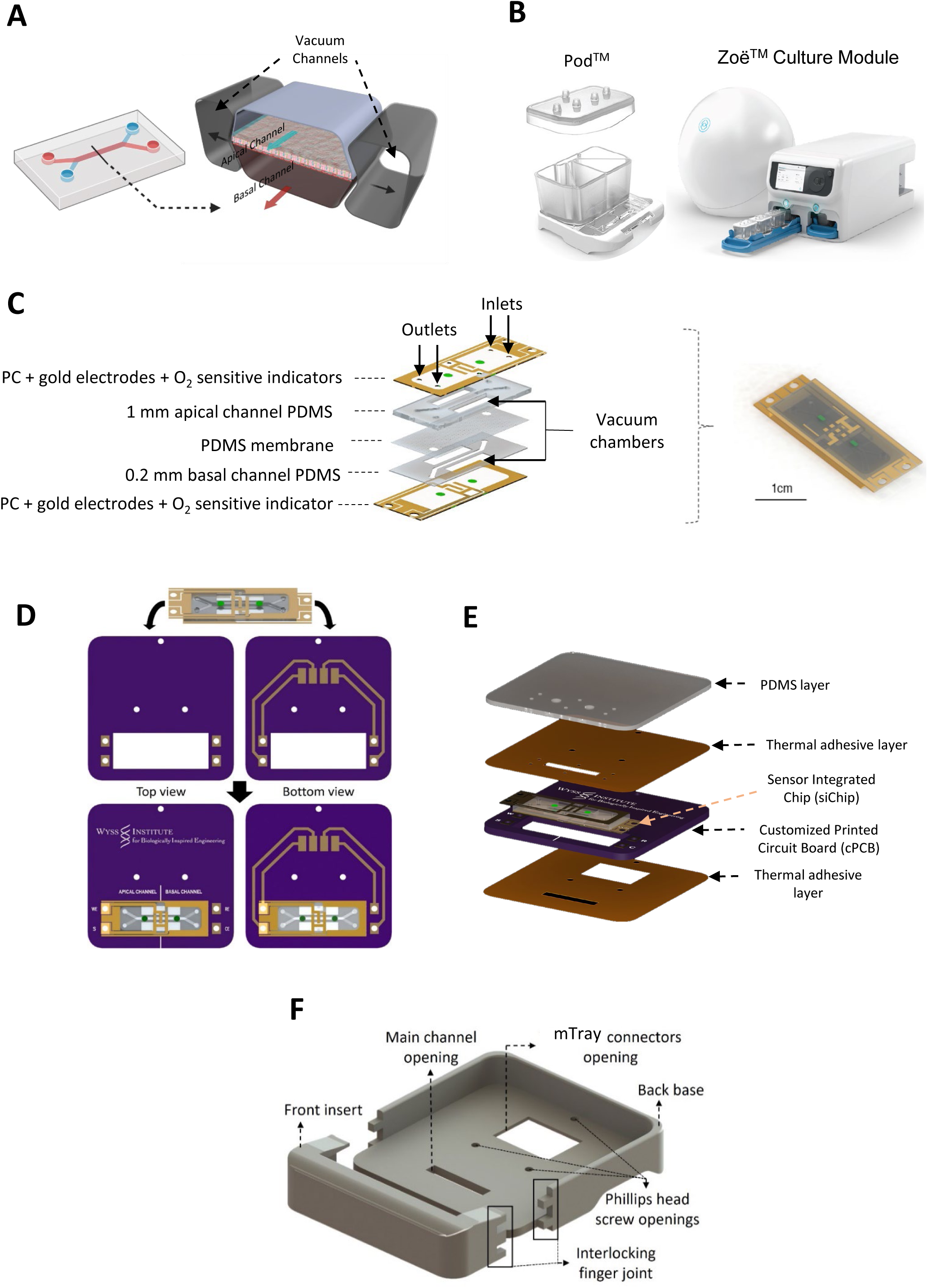
Schematic of commercial organ chip device and culture module and siChip fabrication compartments. **A, B**. Schematic images of the commercial (Emulate Inc, USA) dual channel microfluidic PDMS chip (**A**) and media reservoir Pod^TM^ (left) and Zoë^TM^ Culture Module (right) (**B**). **C**. Schematic image of the layer-by-layer assembly of the siChip made of PC and PDMS layers integrated with TEER and oxygen sensors. **D.** Schematic of the siChip assembly into cPCB. **E.** Layer-by-layer assembly of the siChip to cPCB to make a flushed layer. **F**. Schematic of 3D printed base for assembly of siChip in customized Pod^TM^ (cPod).

**Supplementary Figure. S2.**
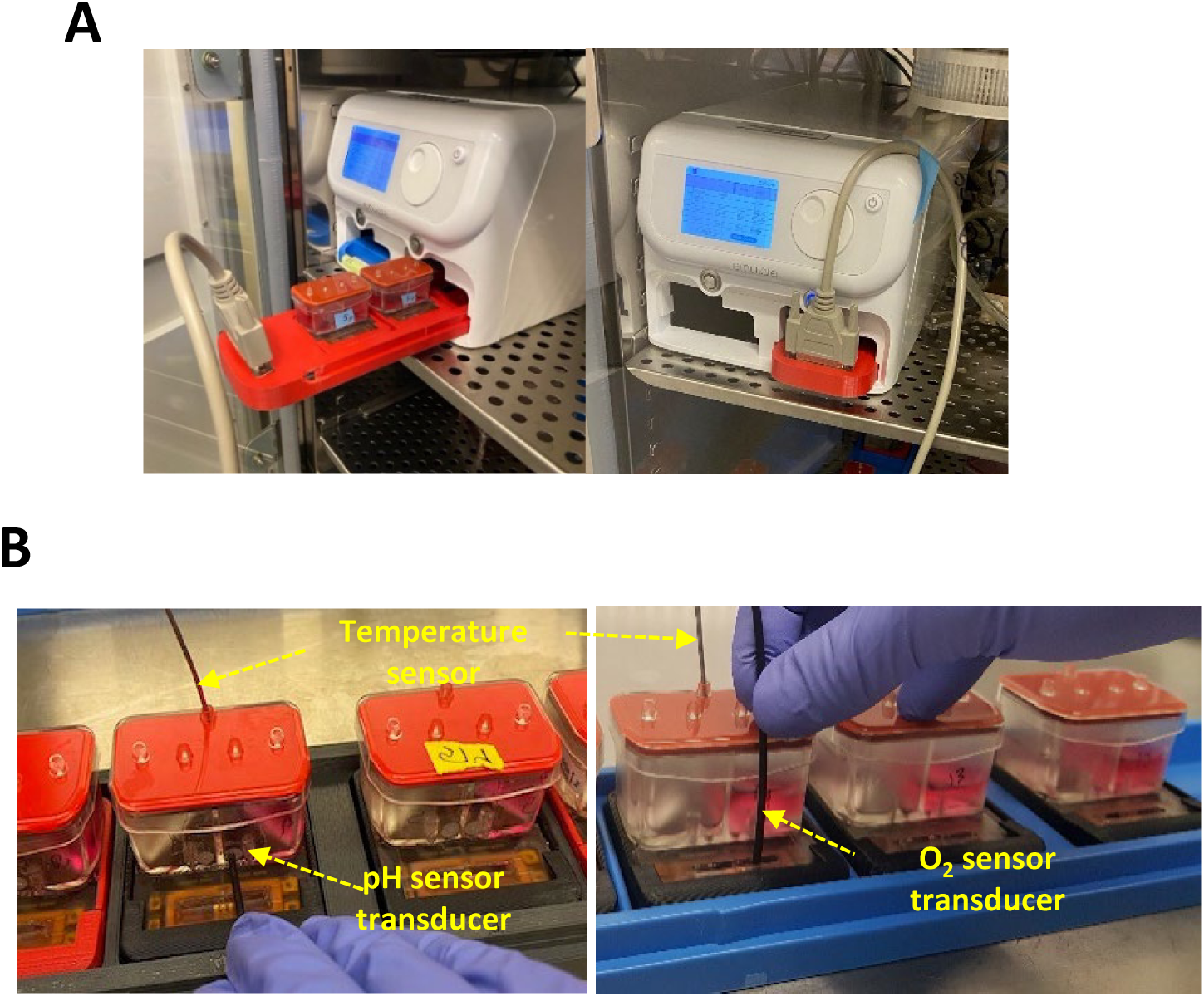
Automated culture and measurement of TEER, oxygen, and pH of siChips. **A.** Photographs of the multiplexed modified Tray (mTray) loaded with siChips for automated culture inside the Zoe culture module (Emulate Inc., USA) with continuous TEER sensing remotely from outside the incubator. **B.** Photographs of non-invasive sensing of temperature, pH (left), and oxygen (right) on siChips.

**Supplementary Figure. S3.**
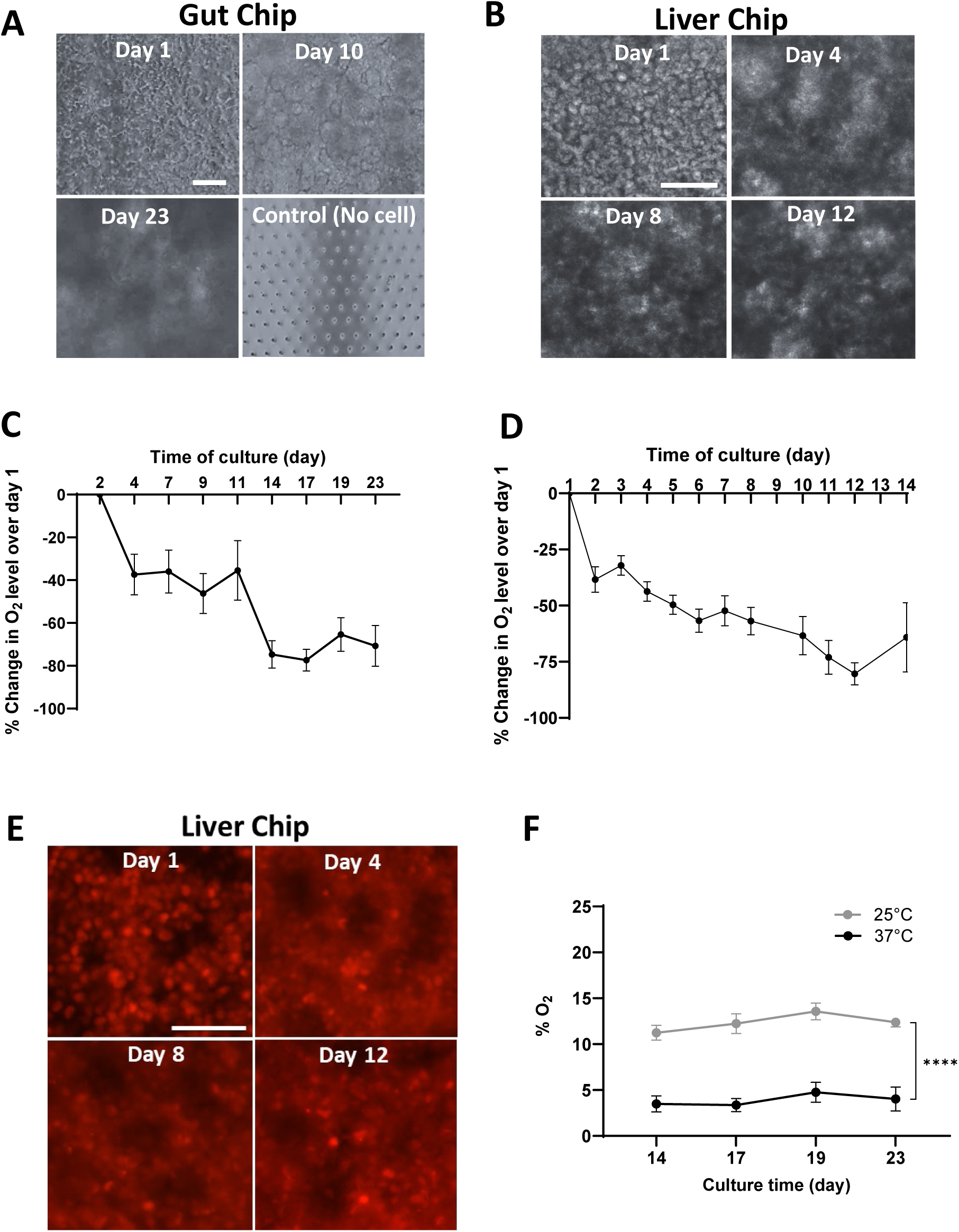
Gut and Liver siChips tissue phenotype and oxygen level dynamics during culture. **A, B.** Phase contrast planar images of Gut (**A**) and Liver (**B**) siChips throughout culture time. Scale bar 100 μm**. C, D.** Continuous monitoring of percentage changes of oxygen level in Gut (**C**) and Liver (**D**) siChips throughout the culture normalized to day 1 or 2 of culture. **E.** Micrograph images of red fluorescence tagged cellular nuclear reporters in epithelium of Liver siChip at day 1-12 of culture. Scale bar 100 μm. **F.** Oxygen levels in Gut siChip at room (25°C) and physiological (37°C) temperatures over 4 days of culture. Data represents the mean ± s.e.m.; n=4-9 (**C**), 5-9 (**D**), and 4-9 (**F**) experimental chip replicate. *\*\*\*\* P< 0.0001*.

**Supplementary Figure. S4.**
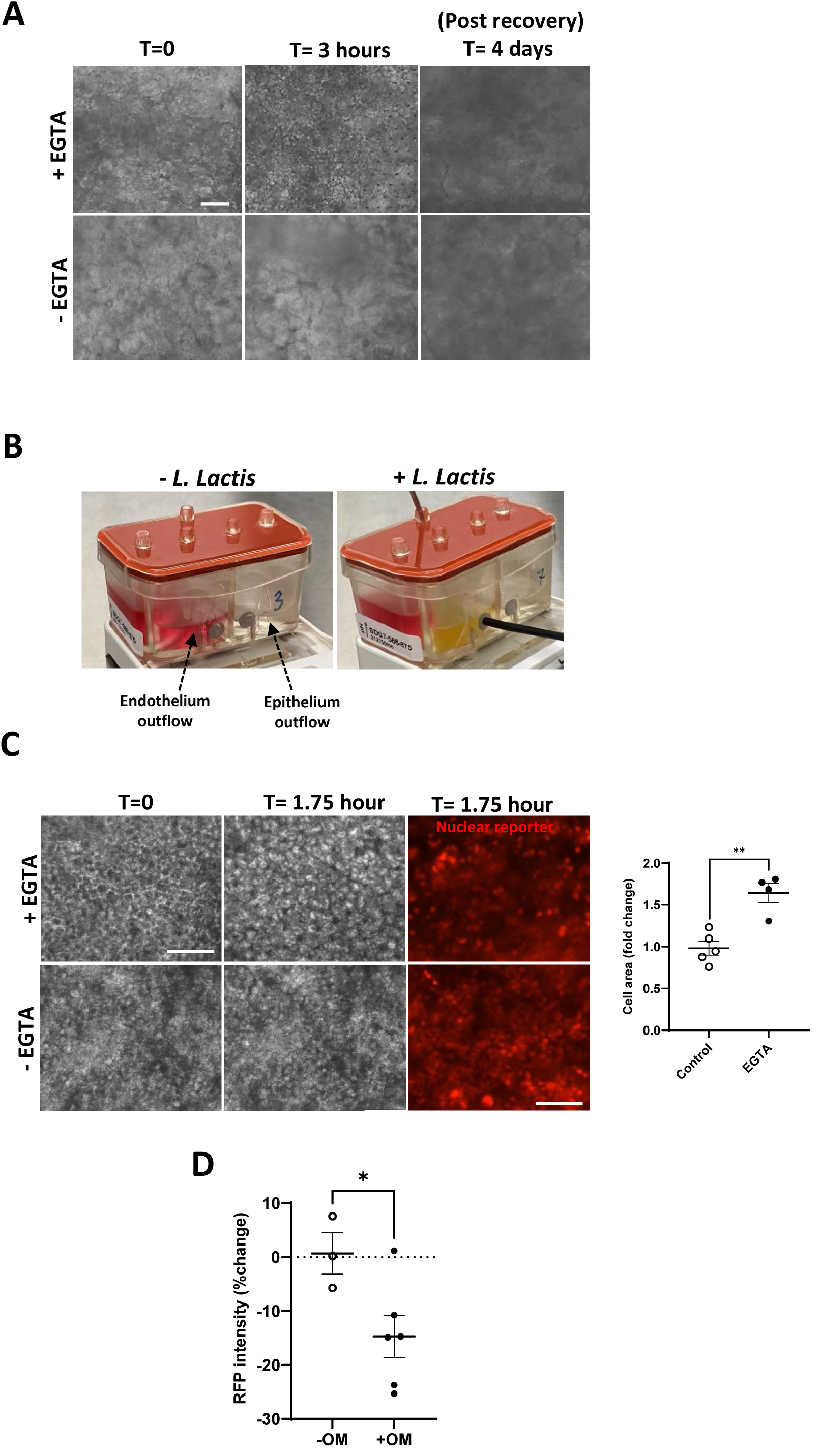
Gut and Liver siChip tissue responses to chemical and bacterial stimuli. **A.** Phase contrast planar images of Gut siChip epithelium before and after exposure to EGTA (+EGTA) and 4 days post termination of treatment compared to control non-treated chip (-EGTA). Scale bar 100 μm. **B.** Photographs of colorimetric pH change in outflow medium of endothelium lumen after 96 hours of co-culture with *L. lactis* bacteria in inoculated chip compared to control chip with no bacteria. **C.** Phase contrast (left and middle) and fluorescent (right) images of liver epithelium before and after 1.75 hours of EGTA treatment (top) compared to control nontreated (bottom) siChips. Scale bar 100 μm. Quantitative analysis of fold change in Huh7 cells surface area after 1.75 hours of exposure compared to control chips (right graph). **D.** Quantitative analysis of percentage change in RFP fluorescent signal intensity of Huh7 cells 3 hours after exposure to OM (+OM) compared to control chips treated with drug carrier (-OM).

